# A Simple Algorithm to Suppress Diagonal Peaks in High-Resolution Homonuclear Chemical Shift Correlation NMR Spectra

**DOI:** 10.1101/2025.08.11.669727

**Authors:** Shengyu Zhang, Jhinuk Saha, Yuchen Li, Xinhua Peng, Ryan P. McGlinchey, Jennifer C. Lee, Ayyalusamy Ramamoorthy, Riqiang Fu

## Abstract

Previous experimental strategies aimed at completely suppressing diagonal peaks in NMR homonuclear correlation spectra often resulted in reduced sensitivity for cross peaks. In this work, we report a spectral shearing approach that transforms diagonal peaks along the diagonal axis of a homonuclear correlation spectrum into a zero-frequency line in the indirect dimension. This allows for effective extraction and substantial suppression of diagonal peaks using a recently proposed data processing algorithm based on quadrature-detected spin-echo diagonal peak suppression. Since the shearing process only rearranges the positions of cross peaks without affecting their intensities, the sensitivity of cross peaks is fully preserved while diagonal peaks are significantly reduced. The effectiveness of this method is demonstrated using uniformly ^13^C&^15^N labeled α-synuclein amyloid fibril and aquaporin Z membrane protein samples.

## Introduction

Solid-state NMR spectroscopy has become an essential technique to determine atomic-resolution structure and dynamics of biomolecules. Among the commonly used techniques is two-dimensional homonuclear chemical shift correlation spectroscopy under magic angle spinning (MAS), which leverages the recoupling of homonuclear dipolar couplings. Such a 2D ^13^C-^13^C homonuclear chemical shift correlation experiment, as shown in Fig. 1a, provides essential long-range carbon-carbon distance restraints for structure determination of ^13^C-labeled biomolecules. However, strong autocorrelated signals, or diagonal peaks, are inevitably present and are typically more intense than the cross peaks. These diagonal peaks can originate not only from the ^13^C-labeled biomolecule under investigation but also from other molecules present in the sample that are not ^13^C-labeled. Since these unlabeled components lack ^13^C-^13^C dipolar couplings, their signals appear only as diagonal peaks. For example, natural-abundance ^13^C signals from lipids present in membrane-mimetics used to reconstitute ^13^C-labeled membrane proteins can produce strong diagonal peaks in the measured 2D ^13^C-^13^C NMR spectrum; and this can be amplified at low temperatures and DNP conditions. Such intense diagonal signals can obscure nearby weak cross peaks, rendering them undetectable. Therefore, it is critical to suppress the diagonal peaks while retaining the cross-peak sensitivity in the homonuclear correlation spectra for the accurate identification and use of cross peaks located close to the diagonal in the structural studies of biomolecules.

**Fig. 1.**
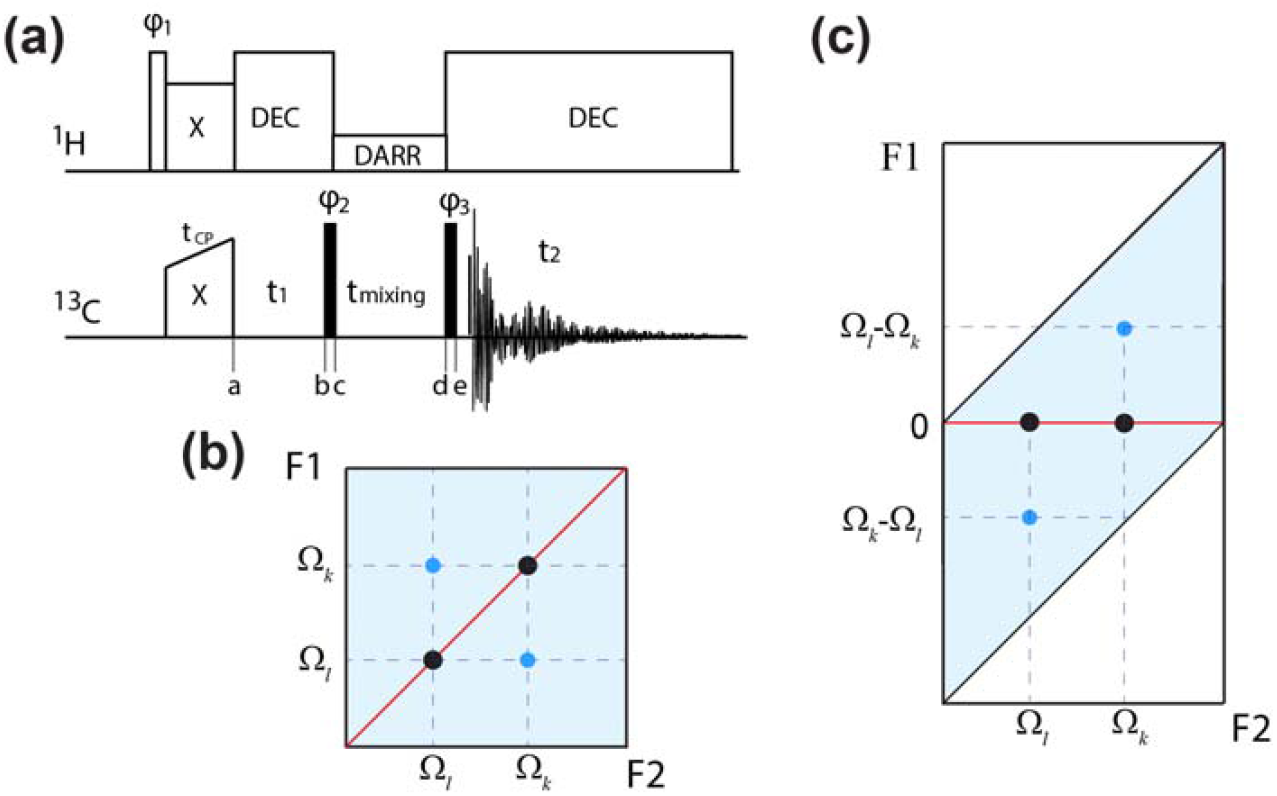
(a) Schematics of the pulse sequence used for standard 2D homonuclear correlation experiments in solid-state NMR under MAS; where DARR, stands for the dipolar assisted rotational resonance irradiation, is used for enhancing the ^13^C-^13^C spin diffusion during the mixing time, and “DEC” represents decoupling irradiation. The open rectangle in ^1^H channel and solid rectangles in ^13^C channel stand for 90° pulses. (b) A 2D spectrum showing the chemical shift correlation for two I and S spins. The red line indicates the diagonal axis. (c) Converted 2D spectrum where the diagonal peaks locate along the zero-frequency line (red line) in the F1 dimension, while the cross peaks locate at their chemical shift difference position.

Double-quantum (DQ) coherence is effective in eliminating signals from uncoupled spins, such as those arising from natural-abundance isotopes or from membrane-mimetic environments and lipids that support membrane proteins [1-4]. For example, 2D DQ-SQ (single-quantum) correlation spectra [5] show only the connectivity between coupled spins, thereby eliminating unwanted background signals from uncoupled spins. However, the efficiency of DQ excitation is generally low, especially for spins that are separated by long distances. As a result, DQ-based methods are limited in their ability to provide long-range distance restraints between labeled ^13^C sites. Therefore, through-space ^13^C-^13^C spin diffusion technique [6-9] remains an effective approach to obtain such long-range distance restraints essential for NMR based structural elucidations of biomolecules.

Subtraction [10-11] between two correlation spectra acquired with different mixing times, one of which mainly generates the diagonal peaks, appears to be a straightforward approach for suppressing the diagonal peaks at the cost of sensitivity and additional experimental time for recording the diagonal peak only spectrum. Recently, Xue et. al [12] cleverly utilized two different signals from cross-polarization (CP) steps: i.e., the CP-transferred signals that participate in spin diffusion process through ^1^H and the non-CP-transferred signals that do not involve in spin diffusion. Subtracting these two signals in every scan leads to partial attenuation of the diagonal peaks with only a limited impact on the sensitivity of the cross peaks. It was demonstrated [12] that the efficiency of the diagonal peak suppression depends on the CP efficiency; thus, a careful intentional CP mismatch setting is required to optimize the diagonal peak suppression at the expense of reducing cross peak intensities. However, this method seems to be successful in 2D ^15^N-^15^N, but not in ^13^C-^13^C, correlation experiments, due to the influence from the proton assisted recoupling effects [13-14]. On the other hand, based on the spin-echo based diagonal peak suppression (DIPS) method [15-16], a sophisticated phase cycling scheme [17] has been used in solid-state MAS NMR to select sine and cosine modulations of the chemical shift difference between the spin-diffused signals in the indirect (i.e., t_1_) dimension, while the autocorrelated peaks appear in the zero-frequency line, which could be effectively suppressed through spectral fittings. However, the cross peaks retain only 50% sensitivity due to the selection of sine and cosine modulations. In addition, the DIPS methods require two t_1_ evolution periods in the t_1_ dimension, as compared to the standard homonuclear chemical shift correlation spectrum, resulting in further decrease in sensitivity due to any unfavored T_2_* relaxation time.

The key component in the quadrature detected DIPS (^QD^DIPS) method [17] is to align all diagonal peaks along the zero-frequency line in the indirect dimension of the resulting spectrum, such that the diagonal peaks can be extracted through spectral fitting and subsequently suppressed from the spectrum. In this study, we report a simple spectral shearing process to transform a standard homonuclear correlation spectrum into the same resonance pattern as in the ^QD^DIPS spectrum where diagonal peaks are aligned along the zero-frequency line in the indirect dimension. As a result, the diagonal peaks can be extracted and subsequently suppressed from the spectrum, rendering a correlation spectrum free of diagonal peaks. Importantly, this is done solely through the data processing and the sensitivity for cross peaks are retained at the same level as that from a regular homonuclear correlation spectrum. Uniformly-^13^C-labeled α-synuclein amyloid fibril and aquaporin Z membrane protein samples are used to demonstrate the effectiveness of this data processing algorithm in terms of diagonal peak suppression.

### Spectral shearing process

Fig. 1a shows the pulse sequence for standard ^13^C-^13^C chemical shift correlation experiments in solid-state NMR. After enhanced through cross-polarization from ^1^H, the ^13^C magnetization evolves under high-power ^1^H decoupling for a period of time t_1_ to express the isotropic chemical shift, followed by a 90° pulse to flip the transverse ^13^C magnetization to the z-axis. Along the z-axis, the ^13^C−^13^C spin diffusion is enhanced through dipolar assisted rotational resonance (DARR) [7, 18] during a mixing time of t_mixing_ to enable the exchange of ^13^C magnetization among different ^13^C nuclei. After the DARR mixing, the z-magnetization is flipped to the xy plane by the second 90° pulse for detection under high-power ^1^H decoupling. To explain the spin dynamics under this pulse sequence and the performance of the algorithm, a simple homonuclear *I*-*S* two-spin system is used to illustrate the correlation between the *k* and *l* spins where their respective chemical shifts are Ω_*k*_ and Ω_*l*_. Assuming no J-coupling is present between these two spins, the observed signals can be represented, when using the quadrature detection in the t_1_ dimension, by:

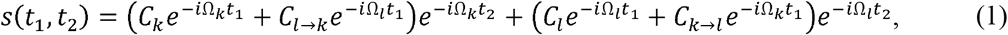

where, the *C*_*k*_ and *C*_*l*_ terms exhibit their own chemical shift modulations in both t_1_ and t_2_ dimensions, corresponding to the autocorrelation, i.e. diagonal peaks, after Fourier transform. On the other hand, the *C*_*k*→*l*_ and *C*_*l*→*k*_ terms carry different chemical shift modulation in the t_1_ and t_2_ dimensions, leading to cross peaks between the *k* and *l* spins, as illustrated in Fig. 1b.

After Fourier transformation in the t_2_ dimension, two 1D spectral slices extracted along Ω_*k*_ and Ω_*l*_ in the F2 dimension of the 2D spectrum can be given as:

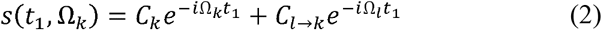

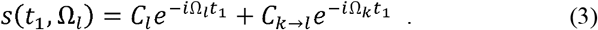

A Fourier transform in the t_1_ dimension, 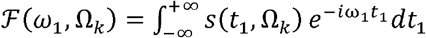 and 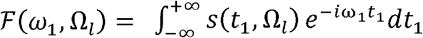, results in a resonance pattern as shown in Fig. 1b.

When applying a given frequency Ω_2_ (i.e., 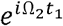) in the t_1_ dimension before the Fourier transformation, we have:

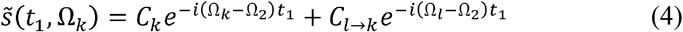

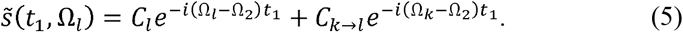

With the Fourier transformation in the t_1_ dimension, we have the following:

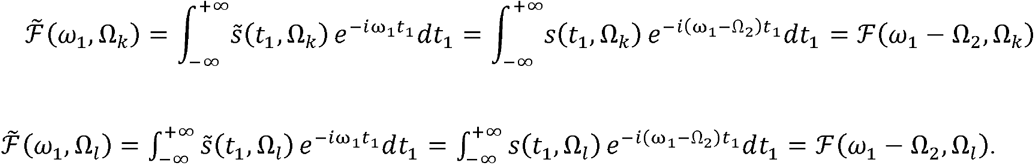

Apparently, this operation simply moves the respective peak positions by −Ω_2_ in the F1 dimension without any impact on the peak intensity: i.e., rearranging the positions from (Ω_*k*_,Ω_*k*_) to (FΩ_*k*_ Ω_2_,Ω_*k*_), (Ω_*l*_,Ω_k_) to (Ω_*l*_ Ω_2_,Ω_*k*_), (Ω_*l*_,Ω_*l*_) to (Ω_*l*_ Ω_2_,Ω_*l*_), and (Ω_*k*_,Ω_*l*_) to (Ω_*k*_ Ω_2_,Ω_*l*_). By stepping this given frequency Ω_2_ through the entire frequency range in the F2 dimension, we can convert Fig. 1b into a new spectrum shown in Fig. 1c. In particular, when

Ω_2_ - Ω_k_ and Ω_2_ - Ω_*l*_, the equations (4) and (5) become (6) and (7), respectively, as given below:

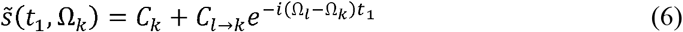

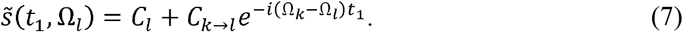

The terms *C*_*k*_ and *C*_*l*_ no longer depend on t_1_, thus representing the spin-echo refocused peaks, while the spin-diffused cross peaks appear at their chemical shift difference positions. In other words, the diagonal peaks along the diagonal axis in Fig. 1b move to the zero-frequency line in the F1 dimension in Fig. 1c. Clearly, the resonance pattern in Fig. 1c is exactly the same as in the ^QD^DIPS scheme[17]. But unlike the ^QD^DIPS scheme where the cross peak intensities are reduced by more than 50% due to the selection of the sine and cosine modulations and using two t_1_ evolution periods, this simple spectral shearing operation does not influence the cross peak intensities at all, as compared to the standard DARR spectra, while the diagonal peaks can be largely suppressed through the same fitting processes as in the ^QD^DIPS scheme [17].

Once the peaks along the zero-frequency line are suppressed, a reverse spectral shearing process (i.e., applying a given frequency 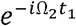 in the t_1_ dimension before the Fourier transformation) can be used to reconstruct the converted 2D spectrum back to the standard correlation spectrum as in Fig. 1b but free of diagonal peaks.

### Experimental

^13^C-^13^C correlation MAS NMR experiments were conducted on a mid-bore 800 MHz NMR spectrometer equipped with a Bruker NEO console, where the ^1^H and ^13^C Larmor frequencies were 799.8 and 201.1 MHz, respectively. Samples were packed into 3.2 mm pencil MAS rotors, and the sample spinning rate was controlled by a Bruker pneumatic MAS III unit at 14 kHz ± 5 Hz. ^13^C magnetization was enhanced via cross-polarization (CP) from ^1^H nuclei using a 1 ms contact time. During CP, a ^1^H spin-lock field of 50 kHz was applied, while the ^13^C B_1_ field was linearly ramped from 38 and 56 kHz [19]. The ^13^C 90^°^ pulse width was 3.0 µs. A SPINAL64 decoupling sequence [20] with a ^1^H B_1_ field of 78 kHz was used during both t_1_ and t_2_ dimensions. State-TPPI method was used for quadrature detection in the t_1_ dimension [21]. A DARR [22-23] mixing period of 50 ms with a ^1^H B_1_ field of 14 kHz was applied during the ^13^C-^13^C magnetization exchange. ^13^C chemical shifts were referenced to the carbonyl carbon resonance of glycine at 178.4 ppm.

MATLAB code was written for the spectral shearing and suppression of diagonal peaks.

## Results and discussion

The processing of the 2D ^13^C-^13^C chemical shift correlation spectrum of α-synuclein fibrils obtained under MAS is illustrated in Supplemental Information Fig. S1. α-synuclein was expressed and purified as reported elsewhere [24]. As expected, the spectral shearing process documented in the previous section moves all peaks along the diagonal axis in Fig. S1a into the zero-frequency line in the converted spectrum of Fig. S1b, while the cross peaks relocate at their chemical shift difference positions. After applying the data processing algorithm developed in the ^QD^DIPS method [17], the peaks along the zero-frequency line are dramatically suppressed as shown in Fig. S1c. The reverse spectral shearing process converts the relocated cross peaks back to their original positions in ^13^C-^13^C chemical shift correlation spectrum while the diagonal peaks are dramatically reduced, as shown in Fig. S1d. For a better comparison, Fig. 2a shows the overlay of the standard DARR spectrum (blue, from Fig. S1a) and its reconstructed correlation spectrum (red, from Fig. S1c). Clearly, the strong diagonal peaks along the diagonal axis in the standard DARR spectrum (blue) are largely suppressed in the reconstructed spectrum, while all the cross peaks away from the diagonal axis have almost the same intensities, such as in the C*α*-C*β* cross peaks at (∼70, ∼60) ppm of the threonine residues. For the methyl rich sidechains at ∼21 ppm, any cross peaks can hardly be identified in the blue standard DARR spectrum in Fig. 2a. However, as indicated in the red spectrum in Fig. 2a, cross peaks between the methyl groups may exist when the diagonal peaks are suppressed. The 1D spectral slices taken from the 2D spectrum in Fig. 2b further confirm the suppression of strong diagonal peaks in the reconstructed spectrum (red) that were present in the DARR spectrum (blue), while with nearly no change to the cross peak intensities. We note that, as indicated in panel II of Fig. 2b, the signal intensities in the red spectrum appear to be less than that in the blue spectrum, even when they are more than 8 ppm away from the diagonal peak position whose linewidth at half height is only 1.5 ppm. This is because the baseline from a strong peak extends far beyond its peak position and can still affect any weak signals in a large range. As an example, the rigid water ^17^O signal is completely buried by the abundant mobile water about 12 ppm away having a linewidth of only 0.7 ppm and can only be observed upon a dramatic suppression of the abundant water signal.[25]

**Fig. 2.**
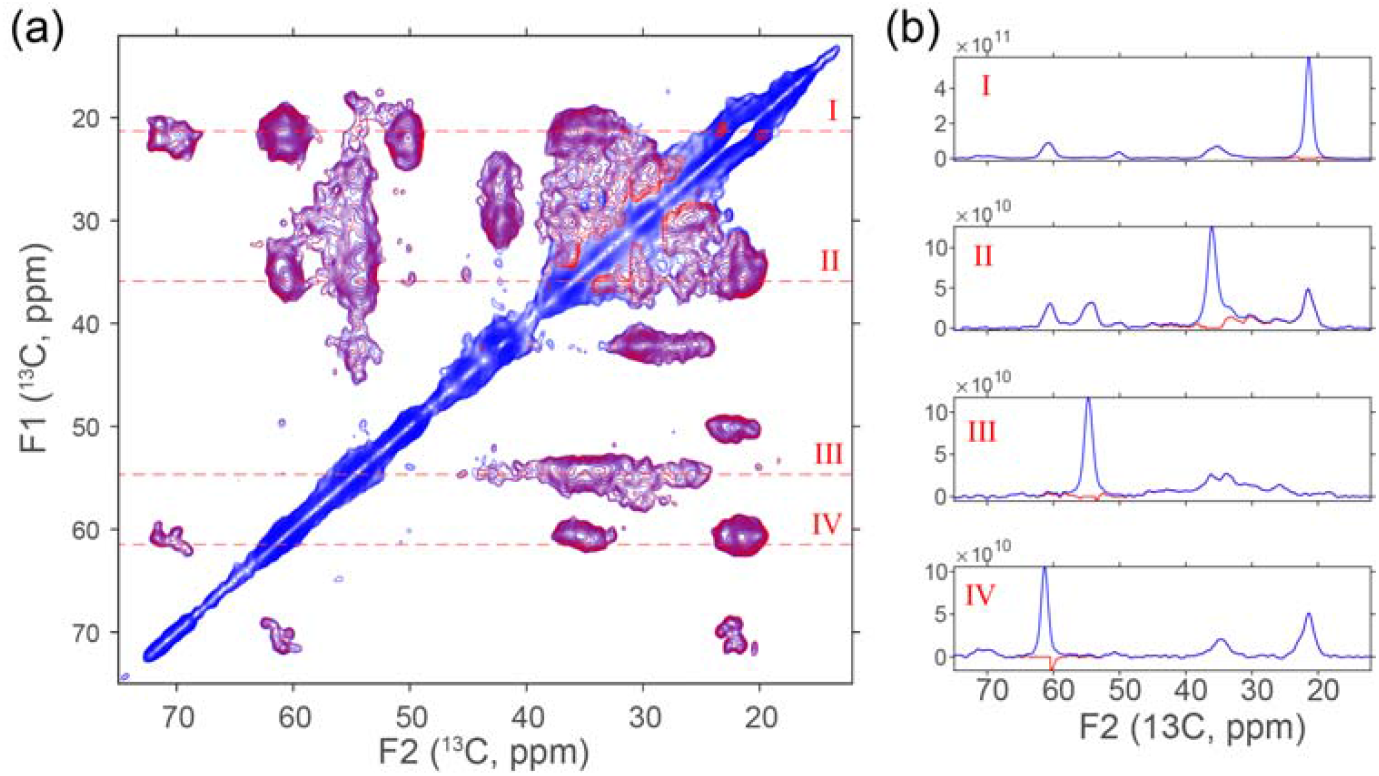
(a) Overlay of 2D ^13^C-^13^C chemical shift correlation spectra of uniformly-^13^C-labeled α-synuclein fibrils obtained under MAS at 275 K: the standard DARR spectrum (blue) and its reconstructed correlation spectrum (red) from the converted 2D spectrum after removing the zero-frequency line. (b) 1D spectral slices taken along the red dashed lines indicated in (a). For the DARR experiment, a total of 512 t_1_ increments was used. For each t_1_, 256 scans coadded each with 2048 FID points and with a recycle delay of 1.5 s. The acquisition times for t_1_ and t_2_ dimensions were 10.24 and 5.12 ms, respectively. The data were zero-filled to a 8192 x 4096 matrix before the Fourier transformation and were processed with a Gaussian window function (LB=-30 and GB=0.1) in both dimensions. α-synuclein was expressed and purified as reported elsewhere. The α-synuclein fibrils were prepared by incubating the monomeric α-synuclein (150□μM) in 10□mM sodium phosphate buffer (pH 7.4) containing 150□mM NaCl and 12□μM DMPG. The mixture was subjected to continuous agitation at 700□rpm at 37□°C for 5 days to promote fibril formation. After incubation, the samples were centrifuged at 17,000□rpm for 30 minutes at 4□°C to pellet the fibrils, and the supernatant containing unaggregated species were carefully removed. Approximately 1□mg of the fibrillar pellet was lyophilized to remove residual moisture and resuspended in 30–35□μL of D□O. The concentrated fibril suspension was then packed into a 3.2□mm MAS NMR rotor for NMR measurements.

Next, we applied this method to the uniformly ^13^C,^15^N labeled Aquaporin Z (AqpZ) in synthetic bilayers. AqpZ is an integral membrane protein that facilitates water across *Escherichia coli* cells with a high rate. The sample preparation for NMR measurements, including protein expression and purification, was detailed in the literature [26]. The processing of the 2D ^13^C-^13^C chemical shift correlation spectrum of the AqpZ protein obtained under MAS is illustrated in Supplemental Information Fig. S2. Similarly, the spectral shearing process documented in the previous section moves all peaks along the diagonal axis in Fig. S2a into the zero-frequency line in the converted spectrum of Fig. S2b, such that the diagonal peaks can be extracted and substantially suppressed using the data processing algorithm developed in the ^QD^DIPS method [17], while the cross peaks relocate at their chemical shift difference positions (c.f. Fig. S2c). The overlay of the standard DARR spectrum (blue, from Fig. S2a) versus its reconstructed correlation spectrum (red, from Fig. S2d) from the converted 2D spectrum after removing the zero-frequency line, is shown in Fig. 3. Again, the diagonal peaks along the diagonal axis in the standard DARR spectrum (blue) are largely suppressed in the reconstructed spectrum (red), while all cross peaks away from the diagonal axis are intact, which are reaffirmed by the slices shown in Fig. 3b. Clearly, the suppression of diagonal peaks leads to the identification of some cross peaks that are close to the diagonal axis in the C*α*-C*α* region as well as in the sidechain region between 20 and 30 ppm.

**Fig. 3.**
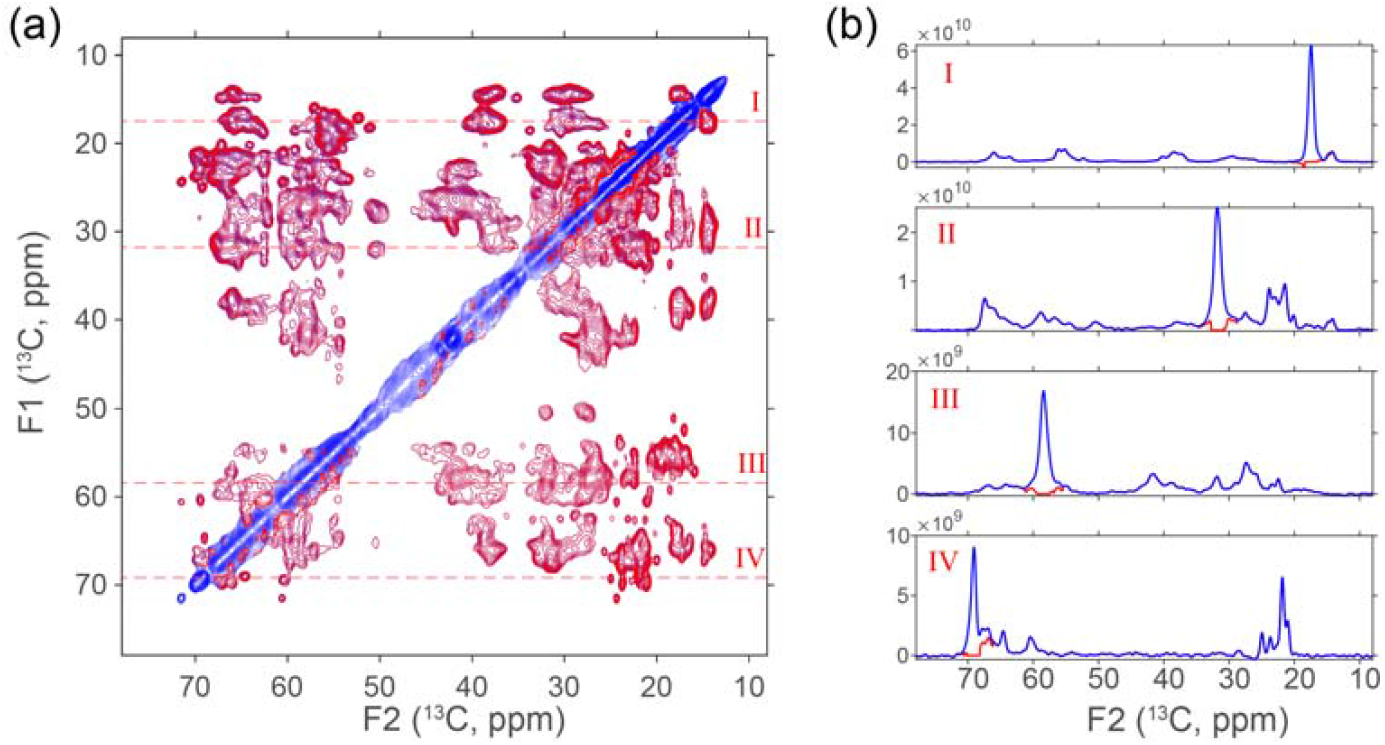
(a) Overlay of the standard DARR spectrum (blue) for Aquaporin Z in synthetic bilayers and its reconstructed correlation spectrum (red) from the converted 2D spectrum after removing the zero-frequency line. (b) Slices taken along the red dashed lines from (a). For the DARR experiment, a total of 600 t_1_ increments was used. For each t_1_, 3072 FID points were recorded at 275 K and 8 scans used for data accumulation with a recycle delay of 1.0 s. The acquisition times for t_1_ and t_2_ dimensions were 15.36 and 6.0 ms, respectively. The data were zero-filled to a 8192 x 4096 matrix before Fourier transform and were processed with a Gaussian window function (LB=-30 and GB=0.1) in both dimensions.

## Conclusion

It has been demonstrated that a simple spectral shearing process converts the diagonal peaks along the diagonal axis in a homonuclear correlation spectrum into the zero-frequency line in the indirect dimension. The diagonal peaks, now aligned along the zero-frequency line, can be effectively suppressed through using a recently proposed data processing algorithm in the quadrature detected spin-echo based diagonal peak suppression [17]. Spectral shearing schemes have been widely used in two-dimensional multiple quantum MAS NMR experiments [27] to correlate the high-resolution isotropic signals in one dimension with their respective 2^nd^-order quadrupolar lineshapes in another dimension. It is well known that a spectral shearing process only rearranges positions of resonances and does not impact their intensities at all. Therefore, once the diagonal peaks along the zero-frequency line are suppressed, a reverse spectral shearing process moves the relocated cross peaks back to their original positions in the homonuclear correlation spectrum without any loss of their intensities, rendering a reconstructed correlation spectrum retaining 100% of the cross peak intensities but free of diagonal peaks along the diagonal axis, as compared to the original correlation spectrum. Importantly, this diagonal peak suppression method is implemented entirely through data processing and can be applied to any processed 2D spectrum. It is thus broadly applicable to various homonuclear systems (e.g., ^13^C, ^15^N, or ^1^H), in both solid-state and solution NMR, including data from published sources.

## Data availability

Data and MATLAB code will be made available upon request.

## Declaration of Competing Interest

The authors declare that they have no known competing financial interests or personal relationships that could have appeared to influence the work reported in this paper.

## Acknowledgements

All NMR experiments were carried out at the National High Magnetic Field Lab (NHMFL) supported by the NSF Cooperative Agreement DMR-2128556 and the State of Florida. We thank Dr. Huayong Xie and Prof. Jun Yang from Wuhan Institute of Physics and Mathematics/Innovation Academy for Precision Measurement Science and Technology, Chinese Academy of Sciences, for providing the AqpZ protein sample. A.R. acknowledges the support from the NIH grant R01DK132214. X.P. acknowledges the support from Innovation Program for Quantum Science and Technology through 2021ZD0303205, National Natural Science Foundation of China through No. 12261160569, and the XPLORER Prize. RPM and JCL are supported by the Intramural Research Program of the NIH, National Heart, Lung, and Blood Institute (ZIA HL001055). LC-MS was performed on instrument maintained by the NHLBI Biochemistry Core.

## Supplementary Materials

Detailed spectral shearing process for the 2D ^13^C-^13^C chemical shift correlation spectra of uniformly-^13^C,^15^N-labeled α-synuclein amyloid fibril and aquaporin Z membrane protein samples.

## Notes

### Competing Interest Statement

The authors have declared no competing interest.

