## supporting information for "A Simple Algorithm to Suppress Diagonal Peaks in High-Resolution Homonuclear Chemical Shift Correlation NMR Spectra"

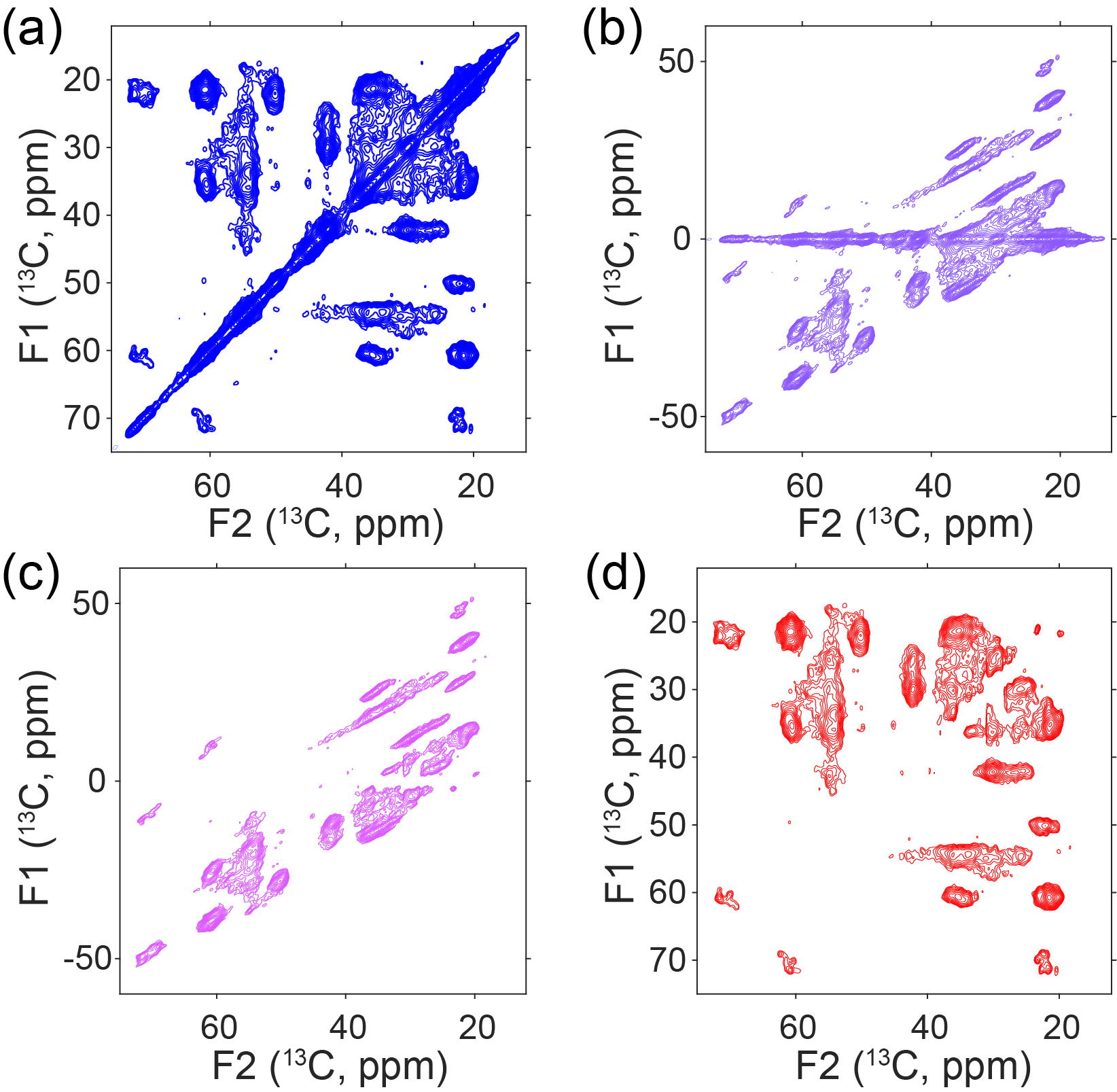

**Fig. S1** Spectral processing of the 2D ^13^C-^13^C chemical shift correlation spectrum of uniformly-^13^C, ^15^N-labeled α-synuclein fibrils obtained under MAS at 275 K. (a) Standard DARR spectrum. (b) Spectrum after spectral shearing, which moves the diagonal peaks from the diagonal axis in Fig. S1(a) to the zero-frequency line in the t_1_ dimension and shifts cross peaks to their corresponding chemical shift difference positions. (c) 2D spectrum after suppression of peaks along the zero-frequency line using the method reported in reference [1]. (d) Reconstructed 2D DARR spectrum with diagonal peaks removed, showing only the cross peaks at their original frequency positions.

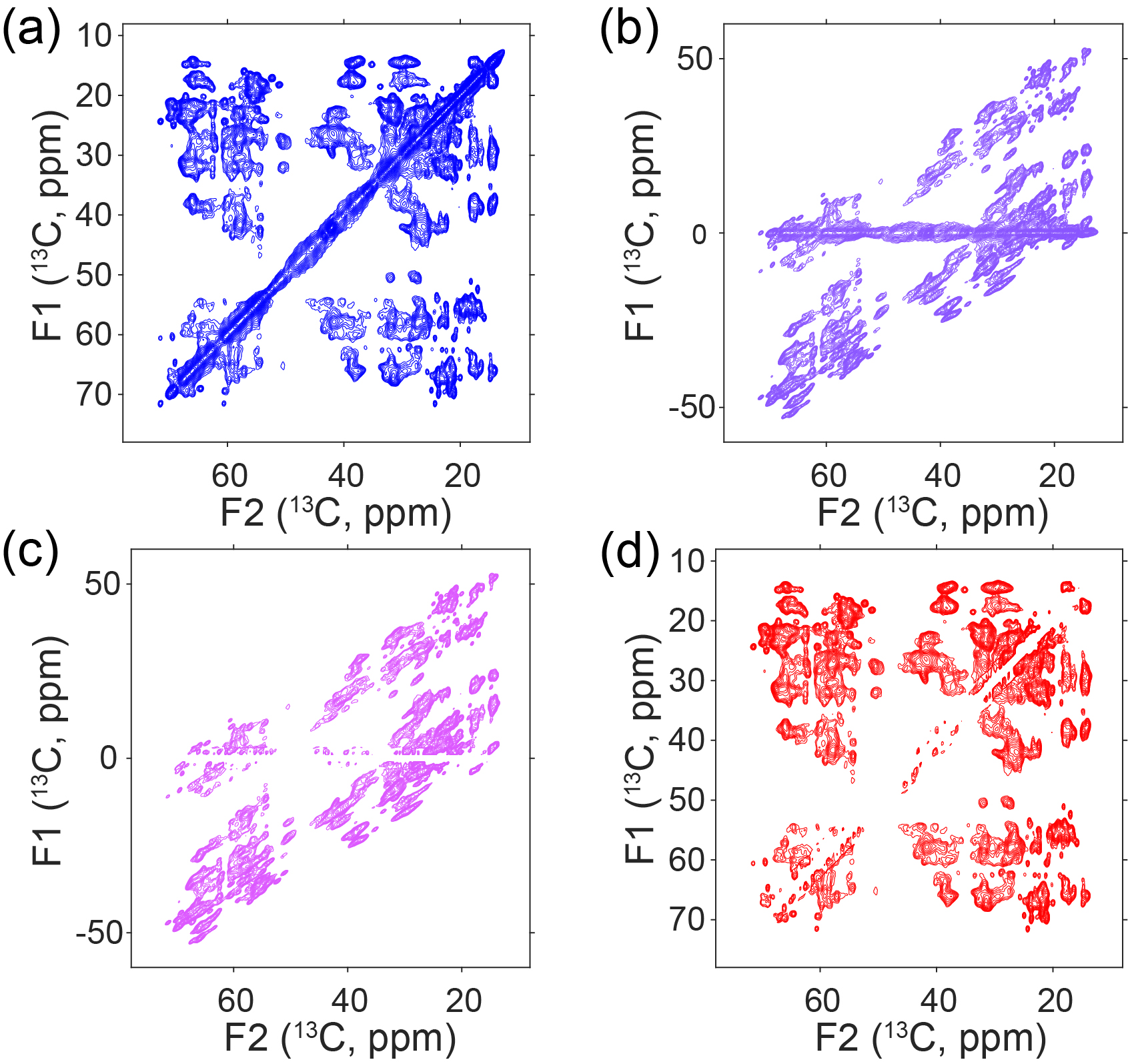

**Fig. S2** Spectral processing of the 2D ^13^C-^13^C chemical shift correlation spectrum of uniformly-^13^C,^15^N-labeled Aquaporin Z protein. (a) Standard DARR spectrum. (b) Spectrum after spectral shearing, which relocates the diagonal peaks from the diagonal axis in Fig. S1(a) to the zero-frequency line in the t_1_ dimension and shifts cross peaks to their corresponding chemical shift difference positions. (c) 2D spectrum after suppressing the peaks along the zero-frequency line using the method reported in the literature [1]. (d) Reconstructed 2D DARR spectrum free of diagonal peaks, showing only cross peaks at their original positions.
